# Characterization of induction methods for *Drosophila* seizure mutations

**DOI:** 10.1101/2021.03.01.433313

**Authors:** Jurga Mituzaite, Rasmus Petersen, Adam Claridge-Chang, Richard A. Baines

**Affiliations:** Division of Neuroscience and Experimental Psychology, School of Biological Sciences, Faculty of Biology, Medicine and Health, University of Manchester, Manchester Academic Health Science Centre, Manchester, M13 9PL, UK; Institute for Molecular and Cell Biology, A*STAR, Singapore; Department of Physiology, National University of Singapore, Singapore; Program in Neuroscience and Behavioral Disorders, Duke-NUS Medical School, Singapore

## Abstract

Epilepsy is one of the most common neurological disorders. Around one third of patients do not respond to current medications. This lack of treatment indicates a need for better understanding of the underlying mechanisms and, importantly, the identification of novel targets for drug manipulation. The fruitfly *Drosophila melanogaster* has a fast reproduction time, powerful genetics, and facilitates large sample sizes, making it a strong model of seizure mechanisms. However, there has not yet been a systematic analysis of the wide range of behavioral and physiological phenotypes observed across major fly seizure genotypes. To understand this, we systematically measured seizure severity and secondary behavioral phenotypes at both the larval and adult stage. Comparison of several seizure-induction methods; specifically electrical, mechanical and heat-induction, show that larval electroshock is the most effective at inducing seizures across a wide range of seizure-prone mutants. Locomotion in adults and larvae was found to be non-predictive of seizure susceptibility. Recording activity in identified larval motor neurons revealed variations in action potential patterns, across different genotypes, but these patterns did not correlate with seizure susceptibility. To conclude, while there is wide variation in mechanical induction, heat induction, and secondary phenotypes, electroshock is the most consistent method of seizure induction across known major seizure genotypes in *Drosophila*.

**Significance Statement:** Epilepsy is a neurological disorder affecting 1 in 130 people globally, with a significant impact on patients, families, and society. Approximately one third of epileptics do not respond to currently available medication. Thus, better insights into underlying disease mechanisms and identification of new drugs are needed. Fruit flies (*Drosophila melanogaster*) are a powerful genetic model: a number of single gene mutant flies exhibit seizures, phenotypes that have been shown to respond to established antiepileptic drugs. We compare methods of seizure induction and their utility, to establish which induction method is the most consistent across a range of different seizure-inducing genetic backgrounds. Adopting a common method for seizure analysis in this model will, we predict, speed identification of novel anti-convulsive treatments.

## Introduction

Epilepsy is one of the most common neurological disorders, affecting ∼60 million people worldwide (Chen et al., 2018). While a variety of causes contribute to epilepsy, including traumatic brain injury and brain infections, the major contribution is from underlying genetic mutations (Poduri and Lowenstein, 2011). It has been estimated that ∼70% of epilepsies do not have a single known cause; of these, 60% have been associated with genetic mutations (Heron et al., 2007; Poduri and Lowenstein, 2011). There are around 700 identified gene mutations currently associated with epilepsy. These include genes contributing to planar cell polarity and the noncanonical WNT signaling pathway (e.g. PRICKLE1), autism spectrum associated genes (e.g. AUTS2) and mTOR signaling pathway genes (e.g. mTOR and TSC1) (Bassuk et al., 2008; Citraro et al., 2016; Wang et al., 2017). However, a majority of epilepsy genes directly influence ion-channel function, specifically mutations in voltage-gated sodium, potassium and calcium channels.

Clinicians have access to over 25 antiepileptic drugs (AEDs) to minimize epileptic seizures (Löscher and Schmidt, 2011). A majority of these drugs target ion channels or neurotransmitter signaling, in an attempt to re-establish an appropriate balance between excitatory and inhibitory signaling in the brain (Löscher and Schmidt, 2011; Vezzani et al., 2011). Although many new AEDs have been approved in recent years, many new drugs have proven no more effective in treating drug-resistant epilepsies than older compounds (Chen et al., 2018; Moshé et al., 2015; Schmidt and Schachter, 2014). It seems likely that treating drug-refractory epilepsy will require novel drug targets and therefore a deeper understanding of epilepsy at the level of basic mechanism(s).

In the genetic model *Drosophila*, both wild-type and mutant animals exhibit seizure-like behaviors; mutants undergo seizures with greatly reduced stimulus thresholds and/or seizure-like activity (SLA) lasts far longer (Baines et al., 2017). In the adult fly, SLA includes repetitive proboscis extension, wing buzzing and loss of posture (Parker et al., 2011a; Tan et al., 2004). These SLAs have formed the basis of a variety of seizure-severity assays (Parker et al., 2011a). In adult flies, seizure-susceptible mutants fall into two main categories based on seizure induction: mechanical (termed Bang-sensitive, BS)- and temperature-induced (Burg and Wu, 2012; Ganetzky and Wu, 1982; Kasbekar et al., 1987; Pavlidis and Tanouye, 1995). Mechanical induction has been standardized with the use of a laboratory vortexer to hyper-stimulate sensory inputs and is termed the ‘vortex assay’ (Kuebler and Tanouye, 2000) (Fig. 1A). By contrast, the heat assay exploits temperature change to induce seizure (Burg and Wu, 2012; Saras and Tanouye, 2016) (Fig. 1B). In larvae, seizures have been induced using a simplified electroshock assay, during which the whole body is subjected to electroshock (Marley and Baines, 2011)(Fig. 1C). A particular advantage of larvae is that they are well-suited to drug screening and, moreover, provide unparalleled understanding of CNS structure and function (Choi et al., 2004; Kadas et al., 2015; Lemon et al., 2015; Marley and Baines, 2011; Worrell and Levine, 2008).

**Figure 1.**
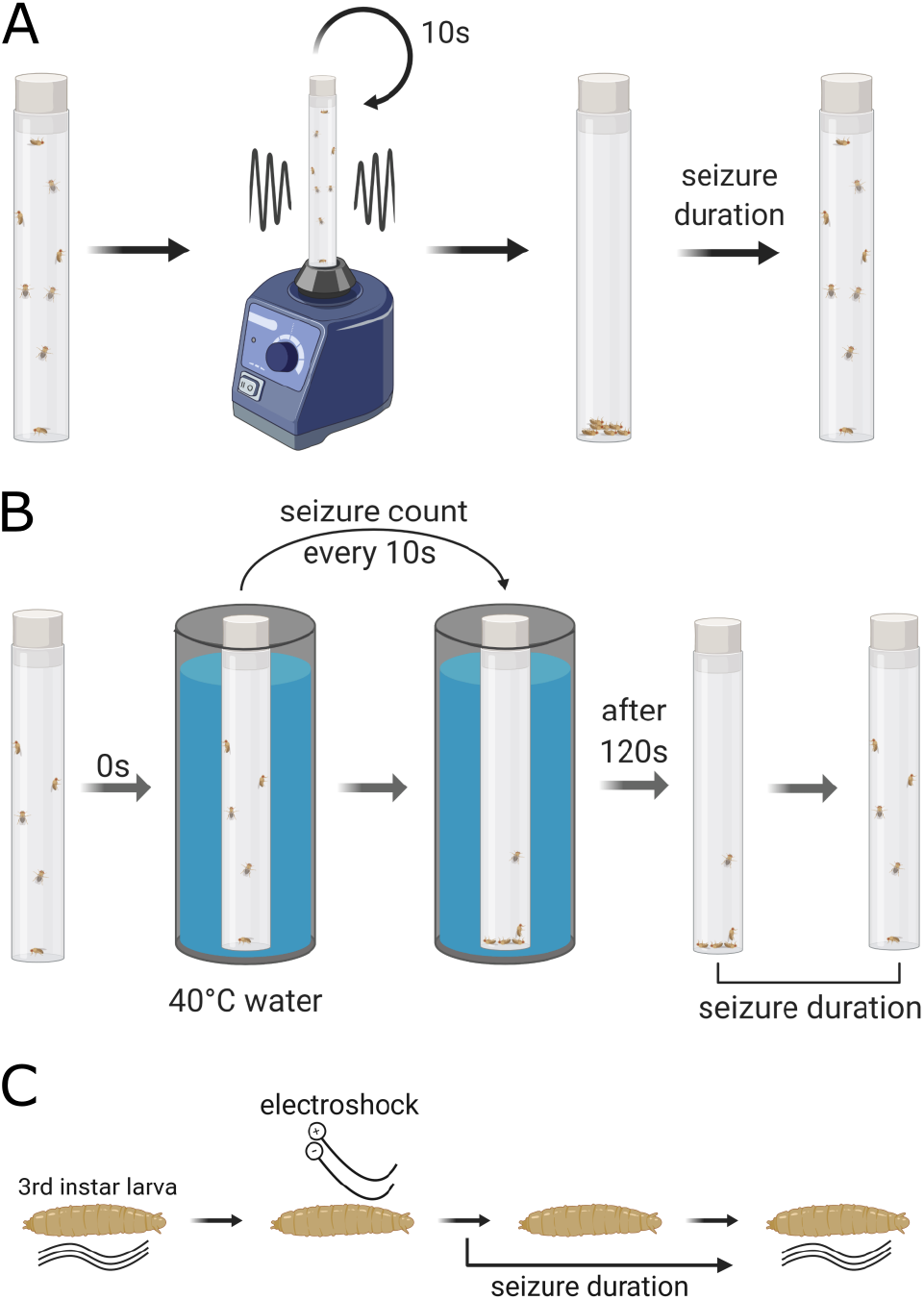
Methods of seizure induction in Drosophila adults and larvae. (A) Schematic showing vortex assay. Adult flies in vials are vortexed for 10s and their seizure duration is measured as time taken to regain their posture. (B) Schematic showing heat assay. Adult flies in vials are exposed to 40°C water for 120s and seizure duration is measured as time taken to regain their posture after being taken out from the water bath. (C) Schematic showing electroshock assay. Electroshock is applied to the dorsal side of 3rd instar larvae. Seizure duration is measured as the time required to restart normal crawling after the shock.

Genetic epilepsies include syndromes characterized by febrile (heat-’fever’-induced) seizures that often present in children. Genetic epilepsy with febrile seizures plus (GEFS+) is commonly caused by sodium channel mutations (Camfield and Camfield, 2015). Some extreme cases of GEFS+ are classified as Dravet Syndrome (DS). This often affects children in their first year of life, and has additional comorbidities including motor and mental impairments (Ziobro et al., 2018). DS is typically resistant to AED treatment necessitating additional research. Use of simpler model organisms, including *Drosophila*, are well suited to investigate mechanisms of epilepsy, but also to identify novel treatments for drug-refractory epilepsies (Griffin et al., 2017, Schutte et al., 2014).

We aimed to establish which accessible seizure-induction methods are applicable across the known range of *Drosophila* seizure mutants. We found that electroshock applied to larvae is the most reliable method to induce seizures regardless of mutation. Since epilepsy is associated with many secondary phenotypes, we investigated if locomotor activity is altered in a predictable fashion, which could be further explored to use for drug screens. We found that there was no significant trend in the daily adult activity levels across the range of mutants tested, but mechanically-sensitive mutant larvae showed reduced crawling. Since electroshock was the most widely applicable method of seizure induction, we investigated potential common underlying mechanisms predisposing larvae to seizures. We detected elongated bursts of motor neuron firing in all seizure mutants. However, the alteration was not exclusive to seizure mutants and did not have good predictive power for seizure severity.

## Materials and Methods

### Fly stocks

Flies were maintained on standard cornmeal medium at 25°C and a 12h:12h light dark cycle. Fly stocks were obtained from the following sources: DS (S1231R), DS-C (S1231S), GEFS+ (K1270T), GEFS-C (K1270K) from Diane O’Dowd (Schutte et al., 2014; Sun et al., 2012); w^-^,eas^2F^;+;+, para^bss1^;+;+, Canton-S and Oregon-R from Richard Baines; w^-^,+;jus^iso7.8^ from M. Tanouye; pk-sple (#422) from the Bloomington Drosophila Stock Centre.

### Adult activity assay

Male flies, aged 4 to 10 days post-eclosion, were placed in 65mm glass tubes containing food in one end and a foam plug at the other. The experiments were carried out at 12:12 hour light-dark cycle in an environment-controlled incubator at 25°C. Fly locomotor activity was monitored for four days, always excluding the first 24 h from data analysis. Animal activity was measured by distance moved per hour (mm/h) and obtained using custom-build image-acquisition software (CRITTA), written in LabView (National Instruments)(Mohammad et al., 2016). All tracking experiments were repeated at least twice, fly N numbers are listed in the respective figures. Prior to tracking, flies were reared at 25°C, collected on the day of eclosion and kept in vials containing only males until the day of experiment.

### Adult vortex assay

Adult flies were collected using CO_2_ 3-4 days post-eclosion into empty plastic vials. 10 flies per vial were left to recover from anesthesia for 1-2h. To assess seizure, the vials were placed on a standard laboratory vortexer at maximum speed for 10s. Seizure duration was measured as time required to regain posture and mobility for each fly.

### Adult heat-shock assay

To measure seizure threshold and duration, adult flies were subjected to a heat assay. Male flies, aged 8–10 days post-eclosion, were anesthetized using CO_2_ and placed into empty plastic fly vials, 5 flies per vial. Flies were given 1–2 h to recover. A water bath was heated to 40–41°C and kept constant at that temperature throughout the experiment. Vials containing flies were placed into the water bath and kept there for 120 s. Throughout the 2 mins in the water bath, visual inspection occurred every 10 s and seizure-status noted. Seizures were identified by loss of posture and random wing buzzing. Vials were taken out of the bath after 120 s and the duration of seizure activity outside the water bath was also recorded (i.e. recovery period).

### Larval locomotion tracking

Larval locomotion was tracked using the DanioVision™Observation Chamber connected to a computer with EthoVision® XT software (Noldus Information Technology, Leesburg, VA). 2% agarose was poured into the lid of 96 well plate and four separate arenas were cut out. Grooves between the arenas were filled with 5M NaCl to prevent larvae from crawling off their respective stages. Individual larvae were placed on the arenas and after a 30 sec adaptation period their locomotion was tracked for 3 minutes. Total distance crawled in the tracking period was calculated by the EthoVision® XT software using centroid tracking.

### Larval electroshock assay

An electroshock assay induces seizure in 3rd instar larvae. Wandering stage larvae were transferred from a vial containing food onto a plastic dish containing water to wash away food residue. Larvae were then transferred to an empty plastic dish and excess water gently removed with a paper towel. No more than four larvae were placed in a single dish. Once normal crawling behavior resumed a probe, composed of two conductive tungsten wires (0.1 mm diameter, ∼1-2 mm apart), was placed on the anterior dorsal surface of a larva, over the approximate location of the CNS. A 7V DC current was applied for 2 s, generated by a DS2A Isolated Voltage Stimulator (Digimeter Ltd, UK). In response to the shock, larvae exhibit sustained contractions of body wall muscles and occasional spasms, halting normal crawling behavior. Seizure duration was measured as a time period between the stimulus onset and resumption of normal crawling behaviour. After seizure, the resumption of normal behaviour involved a full peristaltic wave from either end of the animal which resulted in either forward or backward movement.

### Larval drug treatment

Gravid females and males were kept on grape agar plates (Dutscher, Essex, UK) at 25°C. The flies were fed live yeast paste supplemented by 25 mg/mL picrotoxin (PTX, Sigma-Aldrich, UK) stock to a final concentration of 0.125 mg/mL for three days before embryo collection commenced. Exposed embryos were transferred onto standard food at the end of each collection day and allowed to develop normally.

### Larval electrophysiology dissection

To record from identified aCC or RP2 motor neurons in 3rd instar larvae, the CNS was dissected from a larva and placed in a droplet of external saline solution on a coverslip coated with a thin layer (1-2 mm) of cured SYLGARD Elastomer (Dow-Corning, USA). The external saline solution was composed of 135 mM NaCl, 5 mM KCl, 4 mM MgCl_2_·6H_2_O, 2 mM CaCl_2_·2H_2_O, 5 mM N-Tris [hydroxymethyl]methyl-2-aminoethanesulfonic acid (TES), 36 mM sucrose at pH 7.15. The CNS was stabilized on the coverslip using tissue adhesive (Gluture, World Precision Instruments, USA). The aCC or RP2 cells, located in the dorsal ventral nerve cord (VNC), were exposed for recordings by removing the overlaying glial sheath using a glass pipette (GC100TF-10, Harvard Apparatus, UK) filled with 1% (w/v) bacterial protease type XIV (Sigma-Aldrich, UK).

### Loose patch recordings

Loose patch recordings were performed using borosilicate glass electrodes (GC100TF-10; Harvard Apparatus, UK) pulled to resistance between 1.0 – 1.5 MΩ. The electrodes were filled with external saline of the same composition as listed above. The aCC or RP2 neurons were identified based on location in the CNS, relative soma size and axon configuration (aCC has two axons originating from the soma, RP2 has just one). An electrode was placed on a cell soma without breaking into it. Slight negative pressure was applied to suck around 1/3 of the cell into the electrode. Recordings were carried out using Axopatch 200B amplifier controlled by pCLAMP (version 10.3) using a Digidata 1322A analog-to-digital converter (Molecular Devices, Axon Instruments, USA) or MultiClamp 700B amplifier controlled by WinWCP (version 5.1) using a Digidata 1440A analog-to-digital converter (Molecular Devices, Axon Instruments). Data were sampled at 20kHz and low pass filtered at 0.1kHz. All recordings were made at room temperature (18–22°C).

### Electrophysiology trace analysis

Analysis of loose patch traces was performed using a custom script written in MatLab. Spike times were gathered using Clampfit (version 11.1). A burst is an event with a minimum of three spikes occurring within 100 ms from one another. A burst ends when no spike is detected within 100 ms following the last spike. The script provides summary data on mean burst duration and standard deviation, burst frequency, spike count per burst and firing frequency. Data were plotted using a custom Python script. MatLab script is available upon request.

### Statistical analysis

Data were analyzed with estimation methods using a Python custom script using the DABEST package (Ho et al., 2019). No significance testing was performed, P-values were reported for legacy purposes only. Data are presented following best practices, showing observed values and effect sizes; means are provided with a standard deviation, effect-size errors are displayed as curves with 95% confidence intervals (generated with bootstrapping). Sample sizes (N) are the total number of animals for behavioural experiments or total number of cells for electrophysiology.

## Results

### Mechanical seizure induction

*Drosophila* mutants with increased susceptibility to mechanical shock are known as

‘bang-sensitive’ (Parker et al., 2011a). There are more than a dozen BS single gene mutants (Baines et al., 2017), including: para*^bangsenseless^* (*bss)* a hyperactive Nav channel (Parker et al., 2011b); *easily-shocked (eas)* an ethanolamine kinase (Pavlidis et al., 1994); *julius seizure (jus;* a.k.a. *slamdance)* with unknown protein function (Horne et al., 2017; Zhang et al., 2002); and *prickle*^*spiny legs*^ *(pk-sple)* a protein involved in microtubule polarity (Ehaideb et al., 2014; Tao et al., 2011). Of these mutants, *bss* has the most severe seizure phenotype (Parker et al., 2011b). In our experiments with vortex induction (Fig. 1A), *bss* similarly exhibited the longest-duration seizures (168.4 ± 11.0 s; Fig. 2A). By comparison, *eas* exhibited an average seizure duration of 107.7 ± 8.9 s and *jus* of 73.6 ± 3.9 s. The majority of *pk-sple* did not exhibit severe seizures.

**Figure 2.**
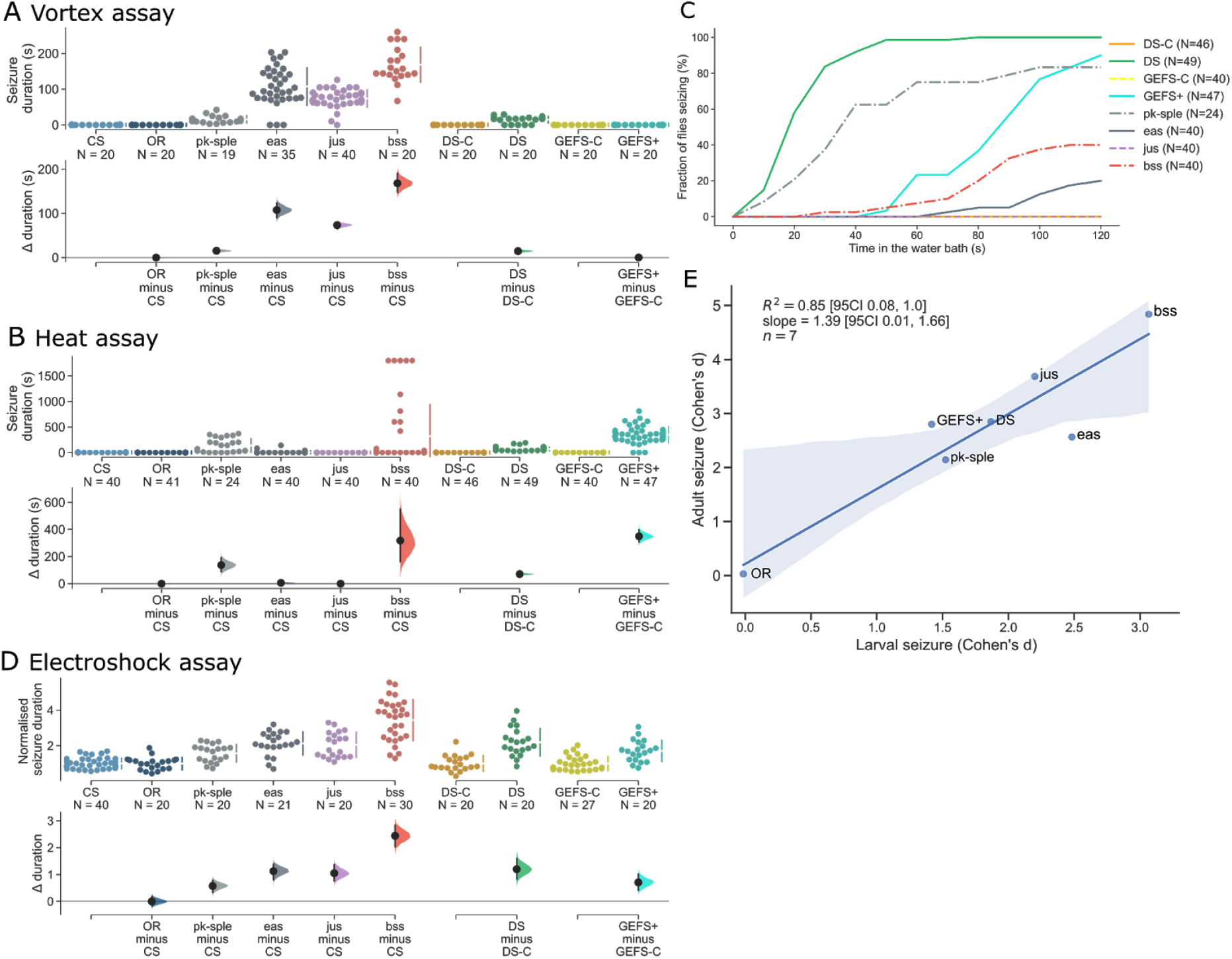
Comparison of seizure induction methods in Drosophila adults and larvae. (A) Vortex-induced seizures in *pk-sple* 15.22 s [95% CI: 11.37, 20.69], *eas* 107.54 s [95% CI: 88.71, 123.84], *jus* 73.35 s [95% CI: 64.60, 79.88] and *bss* 168.35 s [95CI: 157.31, 179.39]. No seizure-like activity was observed in CS, DS-C, DS, GEFS-C and GEFS+ flies. (B) Seizure duration induced by the heat assay. DS showed seizures of average 71.82 s and GEFS+ - 431.03 s, with no seizures recorded for the respective matched controls (DS-C and GEFS-C). *bss* and *pk-sple* showed seizures of 317.33 sec [95% CI: 161.63, 551.63] and 136.67 sec [95% CI: 87.42, 191.84], respectively. By comparison, CS, OR, *jus* or *eas* did not show heat-induced seizures. (C) Cumulative fraction of flies seizing throughout the 120 s of heat-assay. Only 100% of DS flies seized during the 2 min period. Other genotypes reached a maximum of: 90% for GEFS+, 83% - pk-sple, 40% - *bss*, 20% - *eas*. (D) Electroshock induced seizures in all genotypes tested. Results of OR, *jus, eas, bss* and *pk-sple* are reported as a ratio to CS seizure duration which was measured at 97.0 s [95% CI: 95.4, 101.6]. There were no differences between OR and CS. *Pk-sple* exhibited the weakest phenotype with seizure duration increased by 57.47%, whereas *bss* showed an increase of 244.11%. *eas* and *jus* had 112.62% and 104.95% longer seizures. DS and GEFS+ seizure durations are reported as a ratio to their respective controls (DS-C and GEFS-C). DS showed 119.78% and GEFS+ - 70.67% increase in seizures as compared to controls. (E) Seizure induction in larva and adult flies have similar effectiveness (R^2^ = 0.85 [95% CI: 0.03, 1.0], shaded area indicates 95% CI for the linear regression model fit).

Identical stimulation of wild-type lines (CS, OR) resulted in only brief disruption to posture, with a typical recovery of 0.2 ± 0.1 s (Fig. 2A). Both heat-sensitive mutants, GEFS+ or DS, did not exhibit any seizure-like behaviour following mechanical stimulation (Fig. 2A).

### Temperature seizure induction

Several seizure mutants are inducible by elevated temperature (Burg and Wu, 2012; Hill et al., 2019; Kasbekar et al., 1987; Salkoff and Kelly, 1978). Significantly, there are two lines which model specific human epilepsy mutations: GEFS+ (Sun et al., 2012) and DS (Schutte et al., 2014). These ‘humanized knock-in’ mutants exhibit seizures after exposure to elevated temperatures of 38°C and higher (Schutte et al., 2014). We tested the same range of mutants in heat-shock (Fig. 2B, C) as described for mechanical-shock induction, finding that 90% of GEFS+ flies exhibit seizure after 120 s, while 100% of DS flies exhibit seizure after 50 s at 41°C (Fig. 2C). Seizure duration, measured after the heat application was terminated, revealed seizures of average 349 s for GEFS+ and 72s for DS (Fig. 2B). This shows that although DS has a lower threshold for seizure induction, the seizure event is less severe compared to GEFS+.

Some BS lines have been shown to have seizures upon exposure to cold temperatures including *bss* and *eas* at 8°C, but not at 39°C (Burg and Wu, 2012). We find that neither *eas* nor *jus* exhibit seizures in the heat assay (at 41°C, Fig. 2B). Unlike the previous report, we found that *bss* showed seizure behaviour lasting on average 317 s, comparable to seizures in GEFS+ (Fig. 2B). However, only 40% of *bss* flies exhibited seizures, a greatly reduced fraction compared to either GEFS+ or DS. By comparison, 83% of pk-sple tested showed seizure like behaviour with an average time of recovery around 137 sec (Fig. 2B, C).

### Larval electroshock seizure induction

Larvae of *Drosophila* have a peristaltic motor action that has been exploited to study seizure. Application of a brief electric shock to the dorsal cuticle induces SLA (Marley and Baines, 2011) (Fig. 1C). We tested if this seizure-induction method could be used on both the mechanical- and heat-induced seizure mutants tested above. We found that, in response to electroshock, all the mutants exhibited long-lasting seizures (Fig. 2D). Of these, *bss* mutants exhibited the strongest phenotype having seizures 344% longer in duration than wildtype (CS). Additional mutants, *jus and eas,* showed similar duration seizures at 105% and 113% longer than CS, whereas pk-sple seizures were 58% longer. Relative to their own genetic controls, GEFS+ and DS had seizure durations extended by 71% and 120%, respectively (Fig. 2D).

### Seizure induction method comparison

Since electroshock is applied at a different stage of development (i.e. larval) as compared to mechanical and heat-shock (i.e. adult), we compared efficacy of seizure induction methods across the different stages. We used Cohen’s d, a metric for effect size, to compare the efficacy of the vortex assay on BS mutants and heat-assay on HS mutants to efficacy of e-shock in larvae (Fig. 2E). The higher the value of effect size the stronger seizure phenotype was observed in the mutant as compared to control. Although the seizure assay at the adult stage had slightly bigger effect size than at the larval stage, there is a good correlation between the efficacies of seizure induction across the two different stages. This means that seizure severity across the two stages, larvae and adult, is comparable. Thus, electroshock seems to be an efficient and easily applicable method to induce reliable seizures in both mechanical- and heat-induced mutants, making it a favourable method to compare seizure severity across many, if not all, seizure mutations.

### Changes in adult locomotor activity

Alterations in activity patterns and/or sleep duration have been reported for *Drosophila* seizure mutants (Petruccelli et al., 2015). Some epilepsy patients, including DS sufferers present with associated ataxias (Marcián et al., 2016; Oakley et al., 2011). We hypothesized that fly seizure mutants could have similar secondary phenotypes in addition to seizure, and that these may provide ‘simpler’ predictors of seizure susceptibility. To address this, we measured adult locomotor activity over a 3-day period (12:12h cycle) using a custom video-tracking system (Fig. 3A). During the light phase (subjective day) bang-sensitive mutants showed slightly lower activity (Fig. 3B). During darkness (i.e. subjective night) the same seizure-mutant flies tended to have an increased activity compared to wildtype controls (Fig. 3C). The largest difference from CS wildtype during subjective day was exhibited by *eas* at 1012 mm/h reduction, the largest difference during subjective night was also exhibited by *eas,* where it was more active by 667 mm/h.

**Figure 3.**
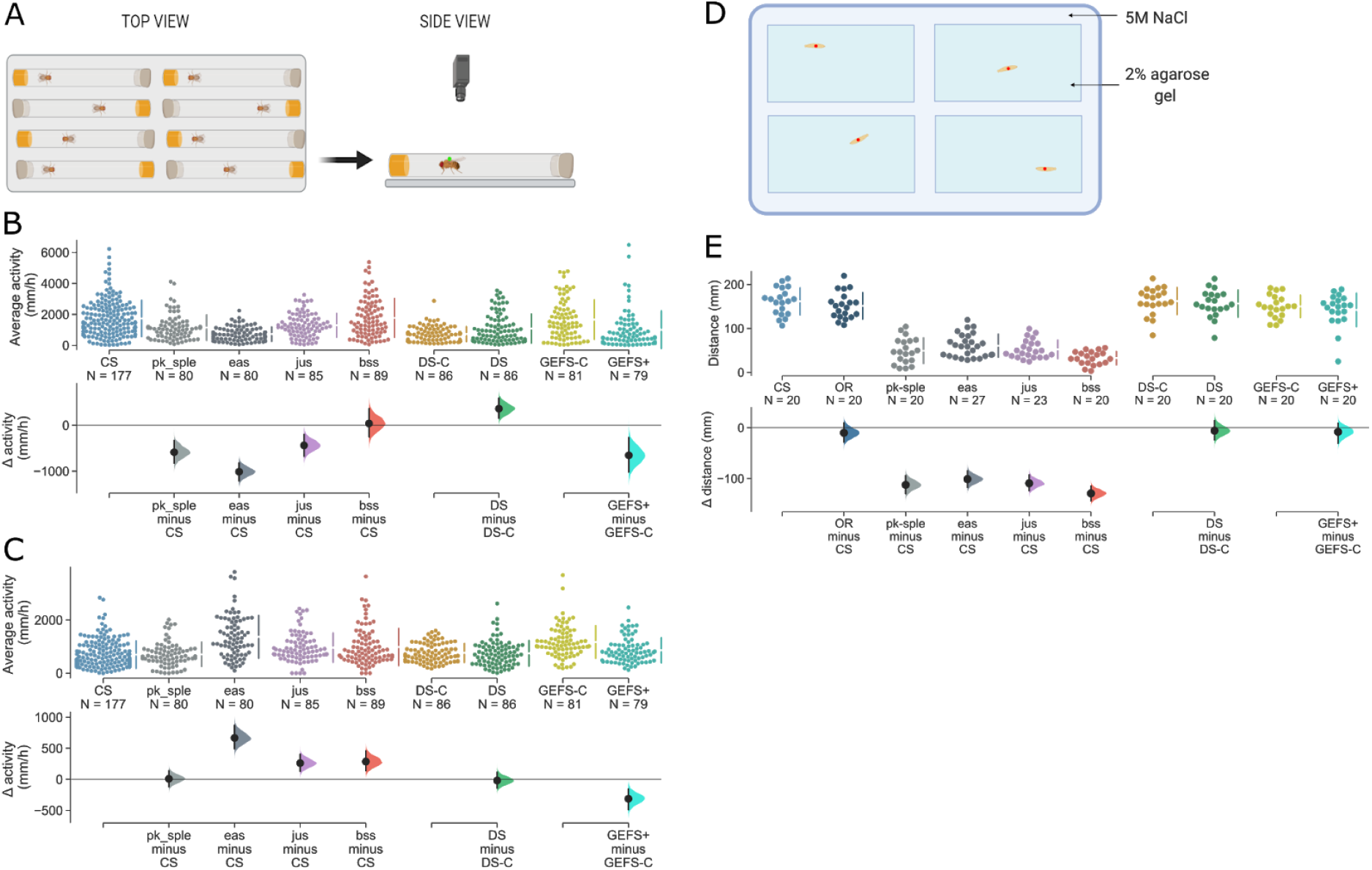
Adult seizure mutants do not exhibit obvious changes in locomotor activity. (A) Schematic showing the locomotor assay. (B) During the 12h period of lights on, *jus* (1298.14 mm/h), *eas* (724.40 mm/h), *pk-sple* (1148.98 mm/h) and GEFS+ (1007.61 mm/h) show reduced locomotor activity (compared to appropriate controls), with *eas* and GEFS+ having the most prominent phenotypes; slower by 1012.09 mm/h [95% CI: −1211.17, −818.82] and 654.49 mm/h [95% CI: −1017.36, −267.84], respectively. DS showed increased activity by 361 mm/h [95% CI: 154.58, 592.13] over its control (DS-C) with an average speed of 727.60 mm/h. (C) During lights off, several mutants showed a small increase in activity levels. Of these, *eas* had the most prominent phenotype at 667.02 mm/h [95% CI: 493.03, 870.27]. *jus* and *bss* showed similar activity levels with an increase of 261.51 mm/h [95% CI: 128.67, 404.60] and 285.95 mm/h [95% CI: 138.50, 458.49], respectively. GEFS+ is the only genotype remaining slower than its respective control (GEFS-C) by 313.14vmm/h [95% CI: −489.98, −154.34]. Other genotypes showed no meaningful differences. Fly activity was tracked for three days at 25°C at 12:12h dark light cycle. All genotypes were recorded at least twice. (D) Schematic showing the set up for larval tracking. (E) Total distances crawled by bang-sensitive mutants *pk-sple* (49.27 mm), *eas* (60.51 mm), *jus* (52.29 mm) and *bss* (32.43 mm) were greatly lower in comparison to CS (161.47 mm). Heat-sensitive mutants did not show any meaningful reduction in the total distance crawled from their matched controls.

Of the heat-inducible lines, GEFS+ was slower by 654 mm/h during the day and by 313 mm/h during night relative to its own matched genetic control. However, DS was 361 mm/h more active in light but showed no difference to the controls in darkness. Out of the lines investigated in this study, DS is the only genotype to have increased activity during the day. GEFS+ was the only genotype to be consistently slower than control.

The stark differences between day- and night-phase activity changes for the differing mutants tested, and the absence of a consistent pattern between activity changes and seizure-induction type or severity, indicate the hypothesis is false: adult locomotor activity is not a reliable predictor of seizure severity.

### Bang-sensitive mutants show reduced larval locomotion

There are previous reports showing alterations in either larval locomotion or peristaltic wave frequency in seizure susceptible *Drosophila* mutants (Graham et al., 2016; Stilwell et al., 2006; Streit et al., 2016). We decided to test for such in the range of mutants used for this study and compare that to the results of adult activity. We performed larval tracking and reported a total distance crawled by each genotype in a three minute period (Fig. 3D, E). CS wildtype had the most activity with a total distance travelled of 161.47 mm. The range of BS mutants showed a consistently lower locomotion level as compared to CS: *pk-sple* slower by 112.20 mm [95% CI:-129.94,-94.07], *eas*- by 100 mm [95% CI:−117.24,-−84.80], *jus* - by 109.18 mm [95% CI:S-124.02, −92.49] and *bss* being the slowest with a reduction of 129.05 mm [95% CI: −143.67, −114.46]. Unlike BS mutants, the range of HS flies did not show any meaningful differences in their locomotor activity among themselves, and in comparison to CS (Fig. 3E). These differences in phenotype reinforce the observation from adult fly activity levels - levels of larval locomotor activity are also not predictive of seizure severity.

### Larval motor neuron activity patterns are not predictive of seizures

Observing that larval electroshock applies to the full panel of seizure mutants we have tested, we decided to investigate electrophysiological properties that may be associated with seizure susceptibility. Because the behavior we observed is due to altered muscular activity, we focused attention to motoneurons. We performed loose patch recordings from the aCC or RP2 motoneurons in third instar larvae. These motoneurons were selected due to their accessibility and prior extensive characterization (Baines and Bate, 1998; Choi et al., 2004). Recordings from aCC/RP2, in wildtype larvae, show a spontaneous, identical and robust bursting pattern of action potentials (APs) which represents the output of the locomotor central pattern generator (Baines et al., 1999) (Fig. 4A). However, alterations in burst recordings were noticed in BS mutants (Fig. 4B). To characterize bursting, we used spike times from recordings of 180 s. We defined a burst as an event with a minimum of three spikes occurring within 100 ms from one another which ends when no spike is detected within the following 100 ms. Based on this classification, recordings from aCC/RP2 from wildtype CS or OR showed one type of bursting pattern (termed ‘normal’) where bursts have an average of 14 or 16 action potentials, respectively (Table 1). Using the same data, we also gathered average burst duration SD, burst frequency, AP count per burst and frequency per burst (Table 1). The data revealed that although there is some variation in all of these parameters, burst duration is consistent with relatively small variation in wild type, but larger variation (in terms of observed standard deviation) in seizure mutants (Table 1). Based on this insight, we classified bursting traces as normal if the average burst duration was the mean of CS burst ± 1.5 SD. Using this classification, we found all mutant lines tested had some normal bursting in addition to some elongated bursting traces (Fig. 4C-E). The fraction of elongated bursting was variable across genotypes, but consistently observed in both BS and HS seizure mutants. Of the mutants tested, *jus* had the highest fraction of elongated bursts among BS (42%, Fig. 4C) and DS among the HS mutants (58%, Fig. 4D). However, unlike the two wild types, DS and GEFS+ matched genetic controls also showed appreciable elongated burst firing. Thus, compared to their respective controls (and not true wild types), DS and GEFS+ showed increased elongated bursting of 17% and 150%, respectively. This, however, suggests that elongated bursting is not exclusive to seizure mutants.

**Figure 4.**
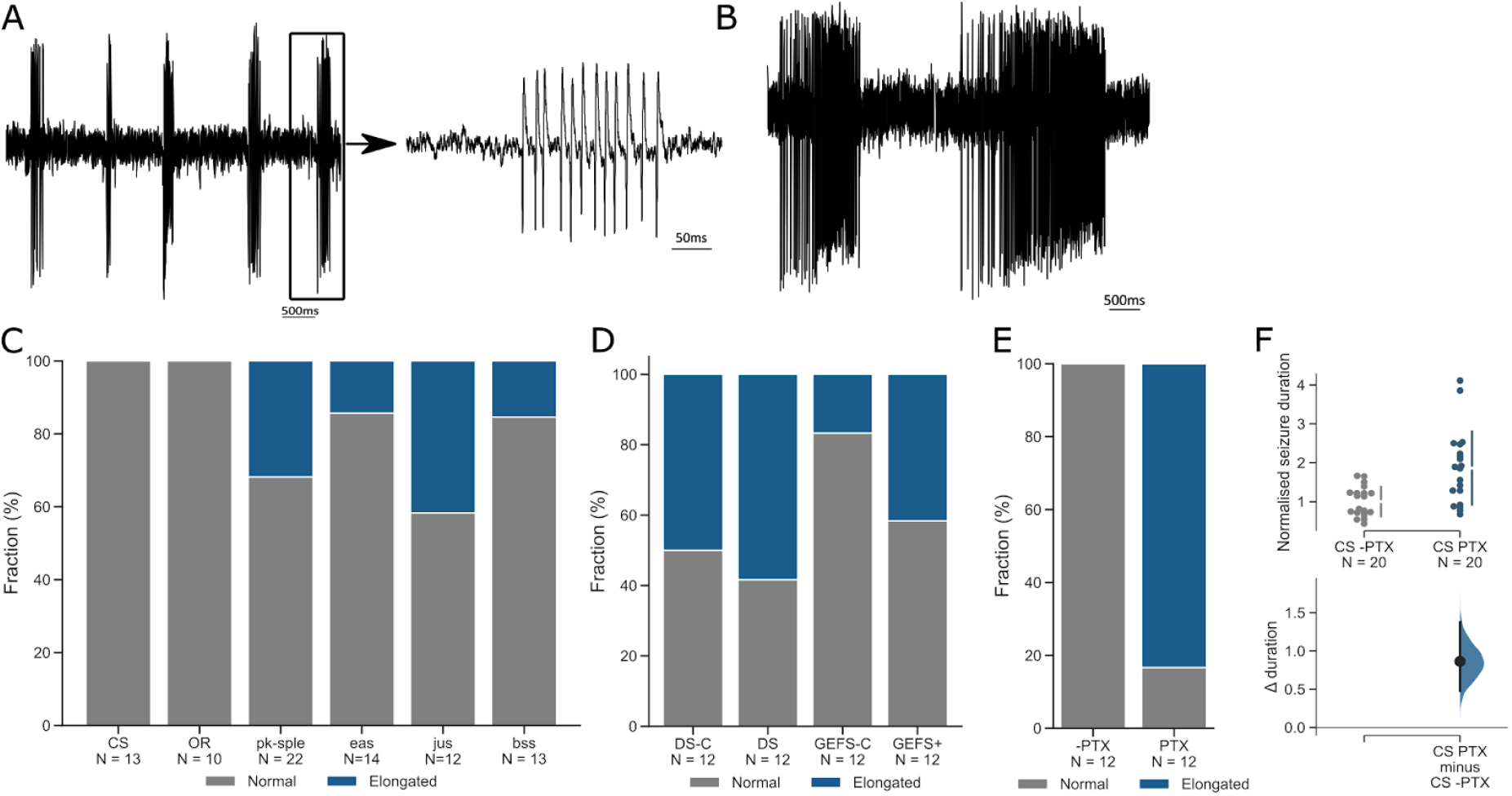
Seizure mutants show elongated motor neuron bursting. (A, B) Examples of cell-attached recordings from larval RP2 and aCC motor neuron bursting patterns in CS wild type (A) and the *bss* seizure mutant (B). (C) The fraction of elongated bursts (i.e. activity shown in B) observed in wildtype and the bang-sensitive genotypes: CS, OR (0%), *pk-sple* (31.8%), *eas* (14.29%), *jus* (41.67%), and *bss* (15.38%). (D) Fraction of elongated bursts recorded in heat-sensitive mutants and their respective controls: GEFS-C (16.67%), GEFS+ (41.67%), DS-C (50.00%), and DS (58.33%). (E) Fraction of elongated bursts detected in CS larvae fed with either vehicle (-PTX) or PTX. Vehicle showed only normal bursting, but recordings from PTX-fed larvae showed 83.33% elongated bursts. (F) PTX induces larval seizures with recovery time 86% longer in the experimental group than control.

**Table 1.** Analysis of activity recordings from larval motor neurons. Three minute loose-patch recordings of spiking activity were analyzed using a custom built MatLab script. Bursts were defined as three or more spikes occurring within 100 ms from each other. The numbers are averages for all traces per genotype.

| Genotype | N | Burst Duration (ms) | Burst Duration SD (ms) | Burst Frequency (Hz) | Firing rate (AP/s) | AP count per burst (AP/burst) | AP frequency per burst (Hz) |
| --- | --- | --- | --- | --- | --- | --- | --- |
| CS | 13 | 174.57 | 91.54 | 0.47 | 6.68 | 14.25 | 87.53 |
| OR | 10 | 155.40 | 84.80 | 0.40 | 6.38 | 16.24 | 104.30 |
| *pk<sup>sple</sup>* | 22 | 394.01 | 348.54 | 0.46 | 6.92 | 16.81 | 51.48 |
| *jus* | 12 | 703.94 | 1211.46 | 0.31 | 8.23 | 23.26 | 58.88 |
| *eas* | 14 | 210.22 | 165.49 | 0.49 | 6.73 | 12.78 | 69.78 |
| *bss* | 13 | 195.42 | 215.28 | 0.43 | 4.97 | 12.72 | 77.89 |
| *GEFS-C* | 12 | 208.94 | 183.28 | 0.37 | 4.32 | 13.67 | 76.16 |
| *GEFS+* | 12 | 1160.34 | 1827.30 | 0.46 | 9.48 | 49.54 | 362.66 |
| *DS-C* | 12 | 11859.25 | 11869.45 | 0.39 | 6.21 | 166.21 | 322.43 |
| *DS* | 12 | 3139.83 | 4338.58 | 0.23 | 4.39 | 42.26 | 34.13 |
| *CS -PTX* | 12 | 170.16 | 89.59 | 0.30 | 4.07 | 12.59 | 90.34 |
| *CS +PTX* | 12 | 1964.82 | 5369.33 | 0.37 | 13.99 | 65.30 | 40.28 |

Picrotoxin is known to induce SLA in wild type flies. The advantage of this approach is that the genetic background is identical between control and PTX-fed animals. We tested for elongated bursting in a PTX-induced seizure model (CS fed PTX) and found that the fraction of elongated bursting was increased to 83%. By contrast unfed control animals showed no elongated bursts (Fig. 4E). PTX-induced seizures in larvae were verified using the e-shock assay, results of which show an increase in recovery time by 86% (Fig. 4F).

We further investigated how aberrations in bursting recordings might correlate with larval seizure (Fig. 5). We used mean burst duration, burst duration standard deviation, burst frequency, AP firing rate, AP per burst count and frequency. None of these parameters revealed any correlation between them and larval seizure severity (Fig. 5 A-F). However, frequency with which abnormalities occur could have a more significant impact on seizure susceptibility than the absolute values of deviations in burst duration. We investigated if the fraction of elongated bursting could be predictive of seizure duration. We found that there is no strong correlation (R^2^ = 0.08) between the two measures (Fig. 5G). The alterations in larvae development could potentially be predictors of adult phenotypes, however, when we compared adult seizure to bursting patterns, the measures were also not well-correlated (R^2^ = 0.14) (Fig. 5H). Thus, we conclude that whilst altered motor neuron activity is typical for seizure mutants, it is not diagnostic for seizure severity.

**Figure 5.**
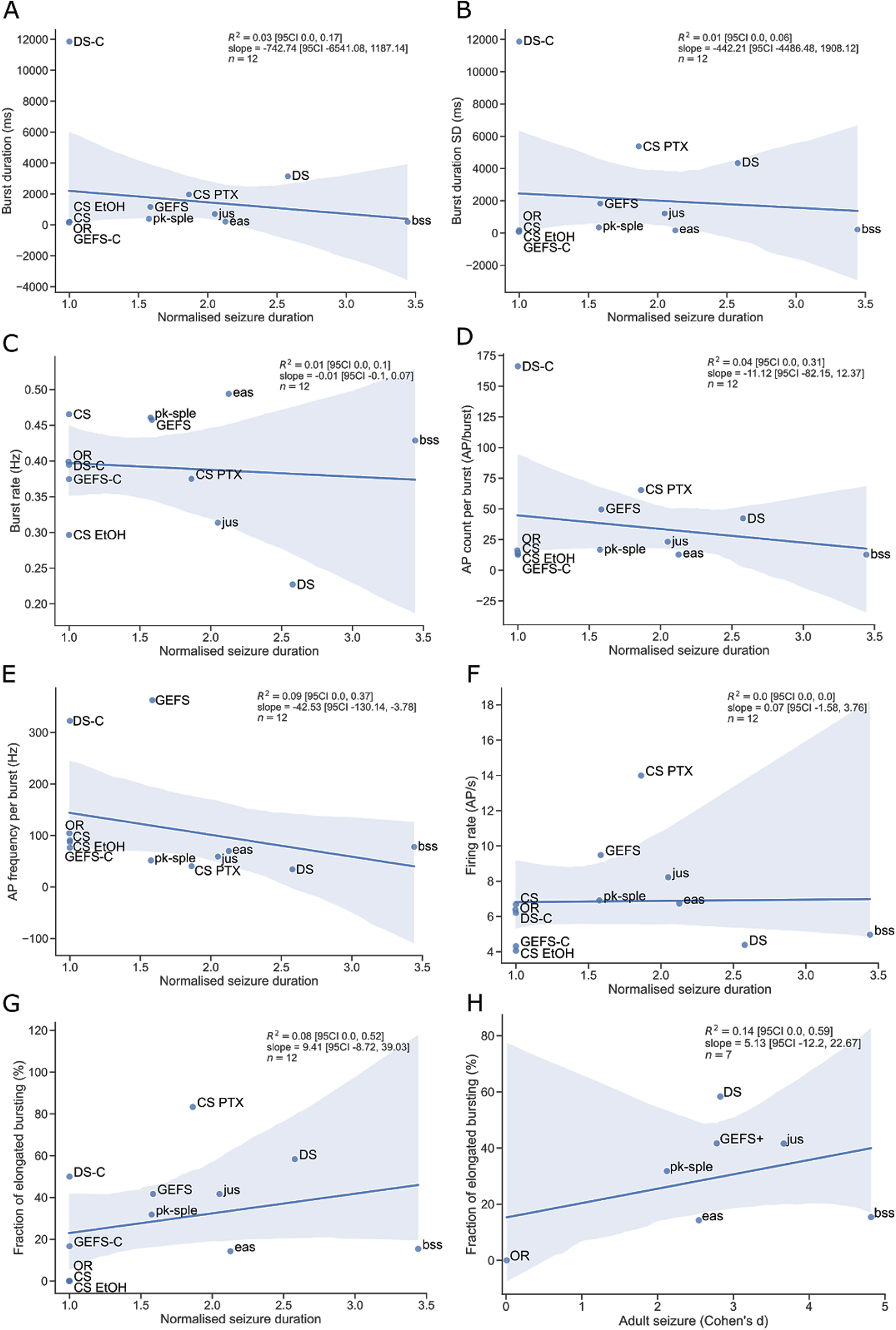
Larval motor neuron firing does not predict seizure susceptibility. Analysis of three minute loose-patch recordings from larval RP2/aCC motor neurons revealed that there is no correlation between mean burst duration (A), standard deviation of burst duration (B), burst frequency (C), action potential (AP) count per burst (D), AP frequency per burst (E), AP firing rate (F) and fraction of elongated bursting (G) and larval seizure. (H) Fraction of elongated bursting from larval motor neurons showed no correlation with the effect size of adult seizure. Shaded areas indicate 95% CI for the linear regression model fit.

## Discussion

Epilepsy is characterized by recurrent seizures, which can manifest as absence seizures, underlying periods of inattention, through to full generalized myoclonic seizures, resulting in muscle jerks to full collapse and unconsciousness. Whilst *Drosophila* seizure mutants lack these full range of behaviours, seizures in both humans and flies nevertheless exhibit sufficient parallels to implicate that the underlying neuronal abnormalities are highly similar. Previous investigations have shown in *Drosophila* that seizures include 1) a defined seizure threshold, 2) genetic mutations that modify seizure-susceptibility, 3) electroshock therapy raises the threshold for subsequent seizures, 4) seizures spread throughout the CNS along defined neuronal tracts, 5) seizures can be localised to distinct regions of the CNS and 6) seizures can be ameliorated by anti-epileptic drugs used to treat human epilepsy (Kuebler et al., 2001; Kuebler and Tanouye, 2000; Lin et al., 2017; Marley and Baines, 2011; Reynolds et al., 2004; Tan et al., 2004). Unlike human epilepsy, however, fly seizure mutants do not routinely exhibit spontaneous seizures which may reflect fewer numbers of neurons within the latter. However, there are a couple of notable exceptions: pk-sple (Ehaideb et al., 2016; Tao et al., 2011) and DS (Schutte et al., 2014). Nonetheless, seizures in flies can be successfully induced using either mechanical, temperature or electroshock induction in a variety of single gene mutant backgrounds, much as seizures can be induced in mammals by extreme external stimuli (light or sound), fever and/or electroshock.

Our study reproduced earlier results validating that all of the lines tested exhibit induced-seizure behavior by a variety of methods (Horne et al., 2017; Parker et al., 2011a; Pavlidis et al., 1994; Schutte et al., 2014; Sun et al., 2012; Tao et al., 2011; Zhang et al., 2002). The length of seizure varies by mutation, with *bss* being the strongest bang-sensitive phenotype in our experiments and, indeed, the strongest reported so far (Parker et al., 2011b). GEFS+ exhibited longer duration seizures than DS following heat-shock, although DS had a lower seizure induction threshold. This is similar to humans, where DS is normally a more severe syndrome than GEFS+ (Brunklaus and Zuberi, 2014).

As in adults, seizures can be induced in *Drosophila* larvae. Electroshock induction has been successfully applied before (Marley and Baines, 2011). These authors reported the *sda* mutant larva (renamed to *jus* (Horne et al., 2017)) exhibit seizure durations up to 6.6 times longer than CS wildtype controls. Likewise, our electroshock induction was effective in inducing seizures in all mutants studied. Seizure duration was variable by genotype, with the longest duration seizures seen in *bss*. Comparing the severity of seizures induced in either adults or larvae suggests that, although adults tend to have longer and more complex seizures, the chosen method for induction is critical. By contrast, our results indicate that electroshock of larvae successfully induced seizures in all mutants and, moreover, generated a comparable hierarchy of seizure severity to adults. We propose that this makes electroshock the most suitable assay for seizure induction in all of the types of mutants studied here and, we would suggest, for studies of novel gene mutations considered to be seizure-related.

We also explored whether adult seizure mutants exhibit pronounced alterations in locomotion, without seizure induction, as has been reported in some human epilepsies (particularly those associated with ataxias). Tracking adult fly activity over several days revealed that there are changes in daytime activity, although these were not consistent across the genotypes studied. During the night, activity was increased; this partly mimics human disorders where epilepsy impairs sleep (Jain and Kothare, 2015; Wang et al., 2018). However, there was again no obvious uniform trend. As such, our results only partially replicated previously observed changes in activity (Petrucelli et al., 2015): whereas the previous study shows an increase in activity during the night, we observed an overall decrease in GEFS+ activity in both day and night. This might be explained by accumulation of genetic modifiers over time. Alterations in food composition are also known to affect seizure phenotype in flies and potential differences in fly food could also impact secondary phenotypes observed (Fogle et al., 2019; Kasuya et al., 2019; Stone et al., 2013). On the other hand, the overall trend we observed matches previously reported data where seizure mutants are, overall, less active than controls (Radlicz et al., 2019). Changes in larval locomotion exhibited a clear difference between mechano- and heat-shock sensitive lines not evident in adult behaviour. BS larvae showed a consistently lowered activity whereas HS larvae did not show any movement phenotype. A previous report shows that both GEFS+ and *bss* exhibit an increase in synchronicity in larval peristaltic waves (Streit et al., 2016), however, in our study it did not translate to the same extent of alterations to locomotion.

Susceptibility to seizures may arise from mutations in a diverse range of genes including those that encode ion channels and/or microtubule polarity-associated genes. However, given a similar behavioral phenotype (i.e. seizure) we explored the possibility of a common underlying change in cellular activity. To investigate this idea, we performed loose-patch recordings from larval motor neurons aCC or RP2 (note, these two neurons are highly similar in their biophysical properties, Baines et al., 1999). In wild type larvae (CS and OR), these neurons exhibit a robust and consistent bursting pattern. The same activity patterns were also present in the seizure mutants and their matched controls. However, in addition, seizure mutants also exhibited altered activity patterns that were not observed in the wild types. This altered activity showed increased burst duration accompanied by modest increases in action potential firing frequency. Increased action potential firing is a hallmark of epileptic seizures (Ehaideb et al., 2016; Marley and Baines, 2011; Parker et al., 2011b). However, our data clearly shows that the presence of elongated burst firing is not well-correlated to induced seizure duration. Thus, at present it is difficult to understand the causes of altered firing activity. The fact that such activity was also observed in both GEFS+ and DS control lines are suggestive that it is not exclusive to seizure mutants. To negate differences in genetic background, that may complicate our results, we fed picrotoxin to CS and observed a greatly increased fraction of elongated bursting. Thus, whilst burst elongation might not be a sufficient explanation for seizure severity, these prolonged activity bursts are clearly associated with seizure *per se*. It is possible that the poor correlation with seizure severity could be due to differing efficiencies of neuronal homeostatic compensation present in the differing seizure mutations we investigated.

In conclusion, *Drosophila* seizure mutants form a diverse group of models for different epilepsy syndromes. Mutants can be separated by seizure induction modes, while larval electroshock is seemingly the most broadly applicable induction method regardless of the mutant genotype. Thus, we recommend this method for testing of novel mutations and its more general adoption might help reduce variability between reported studies that use differing methods of seizure induction.

## Conflict of Interest

The authors report no conflict of interest.

## Funding sources

JM was supported by funding from the University of Manchester, UK and the A*STAR Graduate Academy, Singapore. Work on this project was also supported by funding from Biotechnology and Biological Sciences Research Council (BBSRC, BB/N/014561/1). Work on this project benefited from the Manchester Fly Facility, established through funds from the University of Manchester and the Wellcome Trust (087742/Z/08/Z).

## Acknowledgements

The authors would like to thank Prof Diane O’Dowd (UC Irvine) for providing DS, DS-C, GEFS+ and GEFS-C fly lines and Dr Joses Ho (A*STAR) for help with Python scripts.
Schematics were created with BioRender.com.

## Notes

### Competing Interest Statement

The authors have declared no competing interest.

